# A chemical screen for suppressors of dioxygenase inhibition in a yeast model of familial paraganglioma

**DOI:** 10.1101/2021.11.08.467804

**Authors:** William Beimers, Megan Braun, Kaleb Schwinefus, Keenan Pearson, Brandon Wilbanks, L. James Maher

## Abstract

A fascinating class of familial paraganglioma (PGL) neuroendocrine tumors is driven by loss of the tricarboxylic acid (TCA) cycle enzyme succinate dehydrogenase (SDH) resulting in succinate accumulation as an oncometabolite, and other metabolic derangements. Here we exploit a *S. cerevisiae* yeast model of SDH loss where accumulating succinate, and possibly reactive oxygen species, poison a dioxygenase enzyme required for sulfur scavenging. Using this model we performed a chemical suppression screen for compounds that relieve dioxygenase inhibition. After testing 1280 pharmaceutically-active compounds we identified meclofenoxate HCL, and its hydrolysis product, dimethylaminoethanol (DMAE), as suppressors of dioxygenase intoxication in SDH-loss cells. We show that DMAE acts to alter metabolism so as to normalize the succinate:2-ketoglutarate ratio, improving dioxygenase function. This work raises the possibility that oncometabolite effects might be therapeutically suppressed by drugs that rewire metabolism to reduce the flux of carbon into pathological metabolic pathways.

## INTRODUCTION

Metabolic dysregulation underlies many diseases. Cancer was characterized by a metabolic perturbation now known as the Warburg Effect, the observation that cancer cells exhibit glycolytic, rather than oxidative, metabolism even when oxygen is abundant (Warburg, 1956). Since this discovery, many cancers have been found to have other forms of altered metabolism besides the Warburg effect (Kozal et al., 2021; Pavlova and Thompson, 2016). Familial paraganglioma (PGL) is a remarkable example. PGL is a rare neuroendocrine tumor affecting between 1:100,000 and 1:300,000 people (Berends et al., 2018; Erickson et al., 2001). The most common form of PGL arises in the chromaffin cells that make up the adrenal medulla, where it is known as pheochromocytoma (Lenders et al., 2005). PGL can be characterized hypertension due to secretion of catecholamines, though most cases are asymptomatic. PGL is typically a slow-growing tumor and many cases are benign and curable by surgery. About 25% of PGL cases are hereditary, and most hereditary PGLs are linked to pathogenic variants in nuclear genes encoding the four subunits of the tricarboxylic acid (TCA) cycle enzyme succinate dehydrogenase (SDH; also Complex II of the electron transport chain). Such familial SDH-loss PGL cases thus involve mutations in *SDHA*, *SDHB*, *SDHC*, *SDHD* genes, and in *SDHAF2*, the nuclear gene encoding the factor required for flavin assembly in SDHA. For unknown reasons, variants in *SDHB* are most penetrant (Astuti et al., 2001; Baysal et al., 2000; Burnichon et al., 2010; Hao et al., 2009; Niemann and Müller, 2000). Mutations are inherited as heterozygous loss-of-function alleles, and tumorigenesis is believed to depend upon sporadic mutational loss or silencing of the remaining gene copy in chromaffin cells. Why tumorigenesis is limited to a particular cell type is also unknown.

It is believed that SDH loss of function drives metabolic reprogramming leading to tumorigenesis, but the direct links between SDH loss and transformation are still being established. Current hypotheses focus on the roles of accumulating succinate as an oncometabolite, augmented damage by reactive oxygen species (ROS), and hypersuccinylation (Ishii et al., 2005; Selak et al., 2005; Smestad et al., 2018). ROS production has been shown to increase in SDH-loss model organsims, but it is unclear how much protein and DNA damage results (Adachi et al., 1998; Braun et al., 2019; Ishii et al., 2005; Smith et al., 2007). There is evidence that inherent oxidative stress enhances PGL sensitivity to additional oxidative damage (Liu et al., 2020). Succinate accumulation is perhaps the most promising hypothesis to explain tumorigenesis, with many studies attempting to unravel mechanistic details. SDH loss presumably reprograms central metabolism toward glycolsis because SDH loss breaks the conventional TCA cycle and may alter the flow of high energy electrons into the electron transport chain. Loss of any SDH subunit is believed to disrupt function of the entire SDH complex. This could result in an “obligatory Warburg Effect” with higher dependence on glycolysis, and an accumulation of succinate due to the inability of the nonfunctional SDH to produce fumarate in the TCA cycle (Her and Maher, 2015). However, there is also evidence that chromafin cells may generate ATP from residual steps within the TCA cycle and electron transport chain (Kľučková et al., 2020). Succinate accumulation may extend as well to succinyl-CoA accumulation (one step earlier in the TCA cycle) leading to protein hypersuccinylation (Li et al., 2015; Smestad et al., 2018). The full effects of lysine succinylation on protein function remain incompletely explored. Crucially, accumulated succinate is a competitive inhibitor of an important class of 2-ketoglutarate-dependent dioxygenases. These iron-dependent enzymes oxygenate a substrate by splitting molecular dioxygen. In the process the 2-ketoglutarate (2KG) co-reactant is converted to a succinate byproduct (Loenarz and Schofield, 2011). Accumulated succinate can bind in the enzyme active site, inhibiting such dioxygenases, preventing important chemical transformations in cells (Cervera et al., 2009; Koivunen et al., 2007; Letouzé et al., 2013; Peters et al., 2015; Xiao et al., 2012). A greater understanding of the relationship between succinate accumulation and tumorigenesis is the goal of many studies, but a lack of animal models and PGL cell lines continues to challenge progress.

In an attempt to envision therapeutics and probe PGL mechanisms, several model systems, including *Caenorhabditis elegans*, zebrafish, rodents, mammalian cell lines, and the yeast *Saccharomyces cerevisiae* have been used in lieu of conventional cancer models (Bancos et al., 2013; Braun et al., 2019; Dona et al., 2021; Lussey-Lepoutre et al., 2018; Smestad et al., 2017; Smith et al., 2007). Yeast models of PGL have exploited the conserved mitochondrial role of SDH for studies of metabolic disorders (Bancos et al., 2013; Kregiel, 2012; Lussey-Lepoutre et al., 2018; Smith et al., 2007). *S. cerevisiae* offers many advantages as a model organism, including its fully-sequenced genome and rich genetics (Feyder et al., 2015; Goffeau et al., 1996). The ease of laboratory maintenance and well-documented techniques for yeast analysis make it well-suited for high-throughput assays as well (Bancos et al., 2013). Haploid SDH subunit deletion strains are readily available, and yeast show profound succinate accumulation upon SDH loss, as do human cells (Feyder et al., 2015; Smestad et al., 2018).

Yeast also encode 2-ketoglutarate-dependent dioxygenases, and can therefore serve as a model for succinate accumulation and the subsequent inhibition of these enzymes in this class (Zhang et al., 2020). One dioxygenase in particular, Jlp1p (Fig. 1), is required for sulfur scavenging, converting sulfonates (such as isethionate, ISE) into readily metabolizable sulfites (Hogan et al., 1999). Because Jlp1p is a 2-ketoglutarate-dependent dioxygenase, it can be inhibited by excess succinate. When ISE is the only sulfur source, Jlp1p becomes an essential enzyme for growth. This allows Jlp1p function to be easily monitored simply by assaying cell growth as optical density change over a specified time. In our previous study, we found that a *jlp1*Δ strain is disabled for growth on ISE medium, as expected (Smith et al., 2007). Interestingly, *sdh*Δ strains are also partially disabled for growth on ISE, consistent with Jlp1p inhibition by excess succinate. In contrast *jlp1*Δ and *sdh*Δ strains grow equally well on ammonium sulfate (AS), a sulfur source whose utilization does not require Jlp1p activity (Smith et al., 2007).

**Fig. 1.**
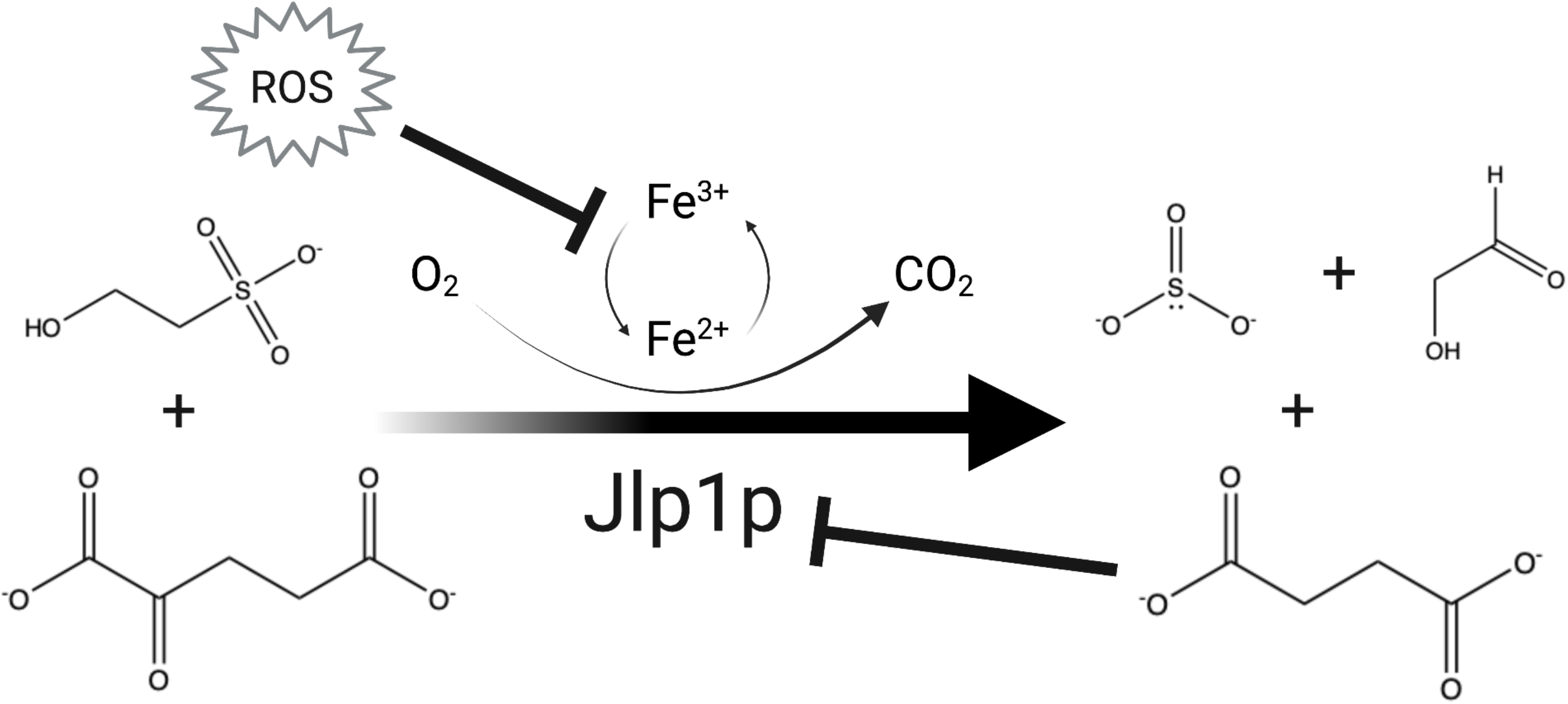
Dioxygenase inhibition in SDH-mutant yeast. Jlp1p dioxygenase conversion of 2KG and sulfonate (ISE) into bioavailable sulfite, with succinate and an aldehyde (glycoaldehyde in this case) as byproducts. The reaction uses molecular oxygen and an enzyme-bound Fe^2+^ ion that undergoes REDOX cycling. In SDH-loss cells, both ROS and succinate accumulate. Depicted are possible mechanisms of Jlp1 inhibition iunder such conditions. Accumulated succinate causes competitive inhibition at the enzyme active site. Increased ROS may interfere with REDOX cycling of Fe^2+^ in the active site.

We reasoned that the dependence of yeast on Jlp1p activity in the presence of ISE as sole sulfur source would provide the potential for a chemical suppression screen to identify compounds capable of mitigating Jlp1p inhibition when SDH is lost. Such compounds might act by reducing succinate accumulation or by moderating ROS production that could inhibit Jlp1p through oxidation of the Fe^2+^ ion required for catalysis (Fig. 1). Based on this concept, we report the results of a screen of the 1280-compound LOPAC library (Sigma #LO1280), and focus on analysis of one particularly interesting compound, meclofenoxate, and its derivative, dimethylaminoethanol (DMAE) as a lead that suppresses Jlp1p poisoning in SDH-loss yeast strains. To the extent that SDH-loss yeast mimic the stresses of SDH-loss human tumors, meclofenoxate and DMAE exemplify a class of compounds that might suppress tumorigenic dysfunction in SDH-loss cells by rewiring metabolism to reduce the flux of carbon into pathological metabolic pathways.

## MATERIALS AND METHODS

### Yeast strains

*S. cerevisiae sdh1*Δ, *sdh2*Δ, *jlp1*Δ strains, and their WT parent BY4742 (MATα *his3Δ1 leu2Δ0 lys2Δ0 ura3Δ0*) were kindly provided by David Katzmann. Strains were maintained on YPGal agar plates (1% yeast extract, 2% peptone, 2% galactose, 2% agar, all w/v), YPGal liquid media (1% yeast extract, 2% peptone, 2% galactose, all w/v) at 25 °C. Most growth experiments were performed using minimal medium supplemented with 20 μM AS or ISE as indicated [Supplemental Table S1;(Cherest and Surdin-Kerjan, 1992)].

### Yeast genotyping

Yeast genotyping was performed as described in supplemental methods by growing a cultures (5 mL) to saturation in YPD (1% yeast extract, 2% peptone, 2% dextrose, all w/v) at 30° C with shaking at 250 rpm.

### LOPAC screen

The Library of 1280 Pharmacologically Active Compounds (Sigma #LO1280) was plated by the Institute for Therapeutics Discovery and Development at the University of Minnesota – Twin Cities. 200 nL of a 10 mM stock of each compound in DMSO were deposited into each well of a 96-well polystyrene plate (Corning #3595). Additional wells received 200 nL of DMSO to serve as controls. For the screen, 200 μL of *sdh1*Δ yeast culture grown to 0.1 OD_600_ in minimal medium containing 20 μM ISE as sulfur source were added to each well (so final compound concentration was 10 μM) and growth was monitored at 600 nm in a plate reader over the course of 24 h at 30°C with constant shaking at 250 rpm. The screen was accomplished with sixteen 96-well plates.

The power of the chemical screen setup was measured using a conventional Z score (see below). Growth of WT yeast in ISE media was compared to the growth of *sdh1*Δ and *sdh2*Δ strains by Z score calculation at multiple time points over the course of 24 h. At each point the Z score was greater than 0, indicating the difference in growth between WT and SDH-loss strains was compatible with a chemical screen. Potential hits were subsequently judged by calculating a Z score according to equation 1:

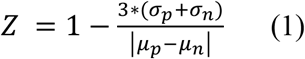

Where *σ_p_* is the standard deviation of experimental growth wells (the *sdh1*Δ yeast strain grown in ISE minimal medium in wells with treatment) and *σ_n_* is the standard deviation of negative control growth wells (the *sdh1*Δ strain grown in ISE minimal medium in wells with vehicle). *μ_p_* and *μ_n_* are the corresponding means of experimental growth wells or negative control growth wells (Zhang et al., 1999). Because each screened compound provided data for a single well vs. 4 data points for the control, the standard deviation of the control was used as a mock parameter for the experimental standard deviation in the calculation. Potential hits were identified from among compounds that stimulated *sdh1*Δ growth with a Z score greater than zero. Compounds formulated with sulfate ions were excluded from consideration as they provided alternative sources of metabolizable sulfur that bypass the reliance on Jlp1p activity.

### Yeast growth assay

WT, *sdh1*Δ, *sdh2*Δ, and *jlp1*Δ strains were grown overnight at 30° C in 10 mL YPGal with shaking at 250 rpm. Cultures were then diluted and grown to mid log phase. Equal numbers of cells of each strain were harvested and washed three times with water by centrifugation at 2500 × *g*. Cells from WT, *sdh1*Δ, *sdh2*Δ, and *jlp1*Δ strains were resuspended in minimal media supplemented with either AS or ISE (20 μM final concentration) to an OD_600_ of 0.1. Yeast culture (100 μL) was pipetted into each well along with 100 μL of vehicle or drug identified as a hit in the LOPAC screen to a starting OD_600_ of 0.05. Each condition was performed with four technical replicates. The growth assay was performed in a 96-well clear plate (Corning #3595) using a SpectraMax Plus 384 UV/Vis cuvette/microplate reader (Molecular Devices; San Jose, CA). The plate was loaded onto the reader and kept at 30 °C without shaking with readings taken every 30 min for 24 h. Aeration was deemed adequate as the oxygen-dependent Jlp1p-catalyzed processing of ISE to sulfite was supported in WT cells. As described, yeast formed a uniform lawn at the bottom of each well for OD_600_ readings (Hung et al., 2018). Growth effect of drug was measured by percent growth difference between treated and untreated at the end of the 24-h growth period. Hits were judged on the magnitude of the effect on *sdh1*Δ and *sdh2*Δ vs WT and *jlp1*Δ. Error was calculated in R through quadruplicate technical replicates using a one-way ANOVA with a post-hoc Tukey HSD test for significance. Growth curves were generated in R and data analyzed using Microsoft Excel and R.

### Metabolite analysis

WT, *sdh1*Δ, and *sdh2*Δ, strains were grown in triplicate for 24 h in 10 mL ISE minimal media cultures supplemented with 100 μM dimethylaminoethanol (DMAE) (Sigma, 471453-100ML). After 24 h, two OD_600_ units of each sample were harvested and media was saved for analysis. Cells were washed by centrifugation twice with PBS, then resuspended in 400 μL H_2_O containing 30 μL concentrated HClO_4_. This cell suspension was vigorously agitated for 25 sec and subjected to three freeze/thaw cycles on dry ice to promote cell lysis. Cell debris was removed by centrifugation and the lysate collected and neutralized with 170 μL 2M KHCO_3_ on ice for analysis as described in supplemental methods.

### Mitochondrial purification and proteomic analysis

WT, *sdh1*Δ, and *sdh2*Δ yeast strains were grown in triplicate to saturation in 1 L cultures at 30 °C with shaking at 250 rpm. Isolation of mitochondria was performed as described from 8 g wet weight yeast (Gregg et al., 2009) and proteomic analysis conducted as described in supplemental methods.

### Western blotting

WT, *sdh1*Δ, and *sdh2*Δ, strains were grown in 10 mL cultures in minimal media supplemented with 20 μM ISE for 24 h at 30 °C with shaking at 250 rpm. Samples were pelleted by centrifugation at 2500 × *g* and washed in DTT buffer (100 mM Tris pH 9.4, 10 mM DTT), then resuspended in 100 μL zymolyase buffer [1 M sorbitol, 20 mM Tris pH 7.5, 50 mM EDTA, 1% β-mercaptoethanol (v/v), 1-2 mg/mL Zymolyase (AMSBIO, 120493-1)] for lysis at 30°C for 15 min. Resulting spheroplasts were pelleted by centrifugation at 2000 × *g* and resuspended in chilled lysis buffer [2% Triton X-100 (v/v), 1% SDS (w/v), 100 mM NaCl, 10 mM Tris pH 8, 1 mM EDTA] and 50 μL chilled glass bead were added. Samples were subjected to 3-5 cycles of vortex mixing (1 min per cycle) with storage on ice between rounds. Samples were subjected to centrifugation at 14,000 × *g* and the supernatant was stored at −80 °C or collected for protein quantification using a BCA assay kit according to the manufacturer’s instructions (ThermoFisher 23227). Equal volumes of whole-cell protein extract were subjected to electrophoresis through denaturing 10% bis-Tris polyacrylamide gels and transferred to PVDF membrane. Equal loading was demonstrated by staining in parallel using Coomassie blue (BioRad). After blocking [5% non-fat dry milk (w/v), Tris-buffered saline, 1% Tween-20 (v/v)], membranes were probed with rabbit polyclonal anti-pansuccinyllysine antibody (PTM Biolabs, PTM-401) at a dilution of 1:1000. Membrane was washed and incubated with IRDye^®^ 800CW goat anti-rabbit IgG secondary antibody (Licor, 926-32211) and imaged using an Amersham Typhoon instrument.

### Fluorescence microscopy

For live-cell detection and quantitation of reactive oxygen species, cells were grown in minimal media (10 mL) at 30 °C with shaking at 250 rpm for 24 h to mid-log phase (0.5 OD_600_). Reactive oxygen species were detected by including H_2_DCF-DA (ThermoFisher #D399) at a final concentration of 10 μM during the 24-h incubation. DHE (Dihydroethidine) ROS detection was performed according to previously published methods (Liao et al., 2020). Cells were then harvested 1 mL at a time, subjected to centrifugation at 10,000 × *g* and washing three times with H_2_O. The resulting cell pellet was resuspended in 20 μL H_2_O, and 5 μL was pipetted onto a glass microscope slide and spread into a thin layer with a glass cover slip. Images were captured at room temperature using an Olympus IX70-S1F2 fluorescence microscope (Center Valley, PA) equipped with an Olympus UPIanApo 100× numerical aperture 1.35 oil objective with the complementing immersion oil (N=1.516, Applied Precision, Issaquah, WA), Standard DeltaVision filters FITC and Rhodamine, and Photometrics CoolSNAP HQ CCD monochrome camera (Teledyne photometrics, Tucson, AZ). Image was acquired using Delta Vision softWoRx (version 3.5.1, Applied Precision, Issaquah, WA) and subsequently processed by FiJi (version: 2.1.0/1.53c, National Institutes of Health, Bethesda, MD). Captured images were exported under the standard DeltaVision file format and converted into 16-bit TIFF images by using Bio-Formats Importer available within Fiji. The contrast and brightness of images were subsequently adjusted within Fiji as well. Data represent quantification from a minimum of three independent labeling experiments with each experiment quantified at least 30 cells. Average cellular fluorescence was quantified using CellProfiler software. Statistical significance was assessed in R by a two-way ANOVA with a post-hoc Tukey HSD test.

### Protein carbonyl colorimetric assay

Protein carbonyl levels in yeast whole cell lysates were assessed using a colorimetric assay (Sigma-Aldrich, MAK094). WT, *sdh1*Δ, and *sdh2*Δ, strains were grown in 10 mL minimal media cultures supplemented with ISE (20 μM) with or without indicated concentrations of dimethylaminoethanol for 24 h at 30 °C with shaking at 250 rpm. Protein lysates were prepared as described for western blotting. Carbonyl levels were assayed according to manufacturer’s instructions. Data analysis was performed in Microsoft Excel. Statistical significance was assessed in R by a two-way ANOVA with a post-hoc Tukey HSD test in R.

## RESULTS AND DISCUSSION

### Yeast strain characterization

As the present study was based on the previous work of Smith et al. (Smith et al., 2007), we began by validating the four experimental yeast strains (WT, *sdh1*Δ, *sdh2*Δ, and *jlp1*Δ) required for interpreting chemical suppression screen results. PCR genotyping confirmed the identity of each strain (supplemental Fig. S1). Growth testing was performed in AS and ISE minimal galactose liquid media to avoid glucose repression while still enabling fermentation and oxidative metabolism (Kayikci and Nielsen, 2015).

Example growth curves for the four strains (Fig. 2 AB) reveal the presence of an expected diauxic shift in the WT and *jlp1*Δ strains in AS medium, indicating sufficient oxygen and an intact TCA cycle and electron transport chain for oxidative growth. In contrast, SDH-loss strains grow only by fermentation. Statistical analysis of growth at 24 h is shown in Fig. 2C. Similar overall levels of growth in AS media are observed, as expected. In contrast, growth in ISE minimal galactose media reveals large differences in fitness, also as expected (Smith et al., 2007). Compared to growth in AS, WT growth in ISE is most robust, with *sdh1*Δ and *sdh2*Δ showing impaired growth, and the *jlp1*Δ strain strongly disabled for growth in ISE media (Fig. 2B), as previously reported (Smith et al., 2007). These growth differences were statistically significant except that growth of the SDH-loss strains was indistinguishable (Fig. 2D). We interpret the growth defect of SDH-loss yeast strains in ISE medium as evidence that Jlp1p dioxygenase function is compromised in these strains, either due to succinate inhibition or oxidative stress affecting the required ferrous ion. These data confirm the findings of Smith et al. and lay the basis for screening for compounds that suppress the *sdh1*Δ and *sdh2*Δ growth defect in ISE media.

**Fig. 2.**
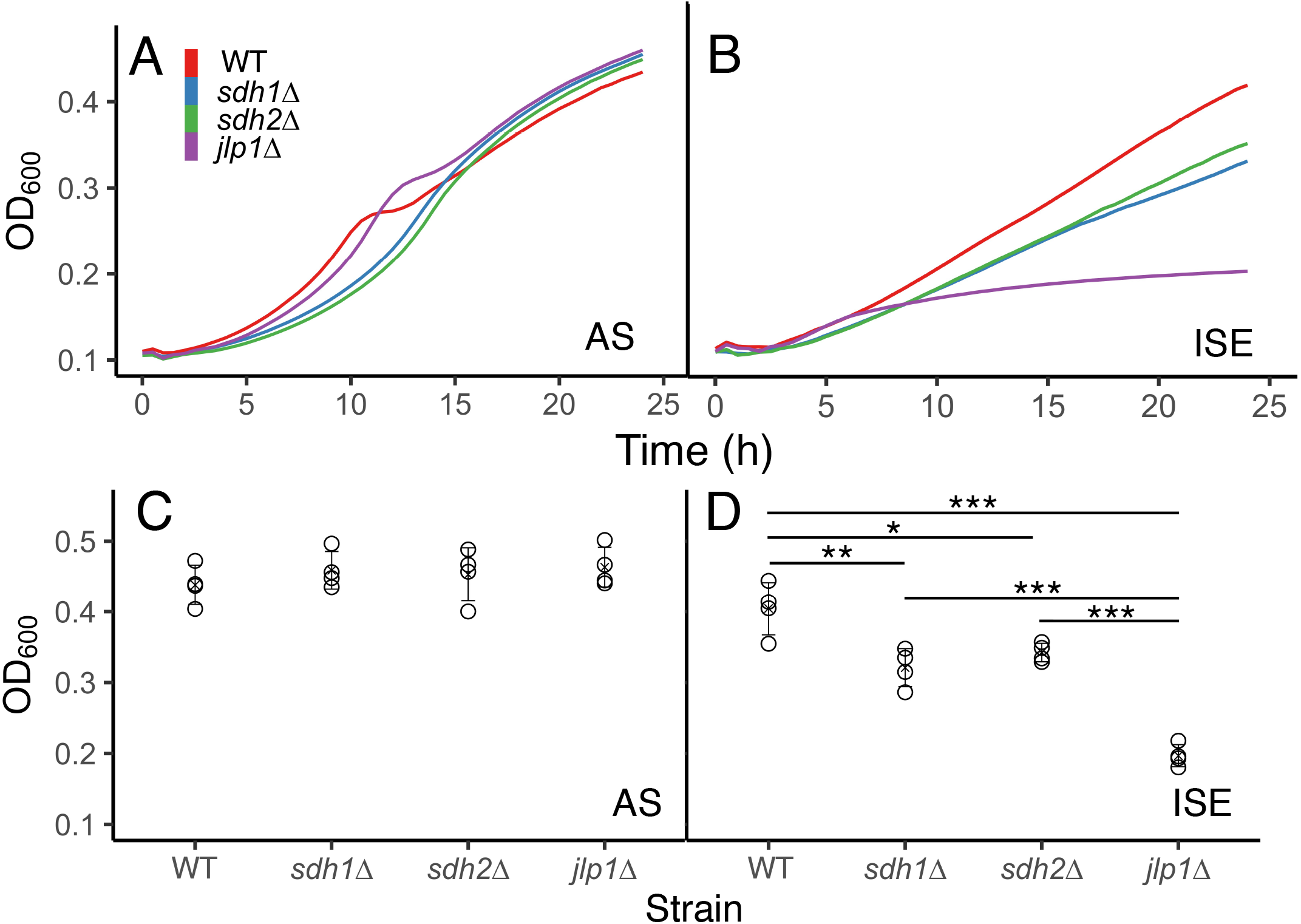
Representative yeast growth phenotypes. A. Growth in ammonium sulfate medium. B. Growth in ISE medium. C. Growth in ammonium sulfate medium at 24 h. D. Growth in ISE medium at 24 h. Statistical significance reflecting four replicates is reported using a one-way ANOVA with a post-hoc Tukey HSD test for significance. P-values: * ≤ 0.05, ** ≤ 0.01, *** ≤ 0.001.

### Proteomic analysis

Previous data indicated that both *sdh1*Δ and *sdh2*Δ strains lack SDH activity but that some Sdh1p protein could be detected in *sdh2*Δ yeast (Smith et al., 2007). In preparation for the chemical suppression screen to identify compounds that rescue Jlp1p activity, we studied potential subtle differences between *sdh1*Δ and *sdh2*Δ strains to select one for screening. We therefore performed a detailed proteomic analysis of WT, *sdh1*Δ, and *sdh2*Δ yeast strains. Mitochondria were isolated from replicate strains grown in YP rich media containing galactose. Extracted proteins were digested with trypsin, peptide lysines acylated using isobaric tags, and the resulting samples analyzed by LC-MS. Approximately 6000 proteins were detected, indicating that even the purified mitochondrial fractions contain a representation of much of the yeast proteome. Quantitative analyses compared detected proteins between WT/*sdh1*Δ, WT/*sdh2*Δ, and *sdh1*Δ/*sdh2*Δ, reported as log_2_(fold change) with statistical significance as an adjusted P-value (significance ≤ 0.05). Volcano plots (Fig. 3AB) indicate differences between WT and SDH-loss strains. In the WT/*sdh1*Δ dataset 1015 proteins were significantly different between WT and *sdh1*Δ yeast, with 911 more abundant in WT and 104 more abundant in *sdh1*Δ. In the WT/*sdh2*Δ dataset 1068 proteins were significantly different between WT and *sdh1*Δ yeast, with 946 more abundant in WT and 122 more abundant in *sdh1*Δ. These results confirm the large remodeling of the yeast proteome driven by SDH loss. We compared the SDH-loss strains in greater detail. Of the 1092 proteins that were differentially expressed upon SDH loss, 991 of these (91%) were shared between both *sdh1*Δ and *sdh2*Δ, 77 (7%) were unique to the WT/*sdh1*Δ dataset, and 24 (2%) were unique to the WT/ *sdh2*Δ dataset (Fig. 3C). This analysis confirms that *sdh1*Δ and *sdh2*Δ are proteomically comparable, and this is affirmed by a correlation plot (Fig. 3D).

**Fig. 3.**
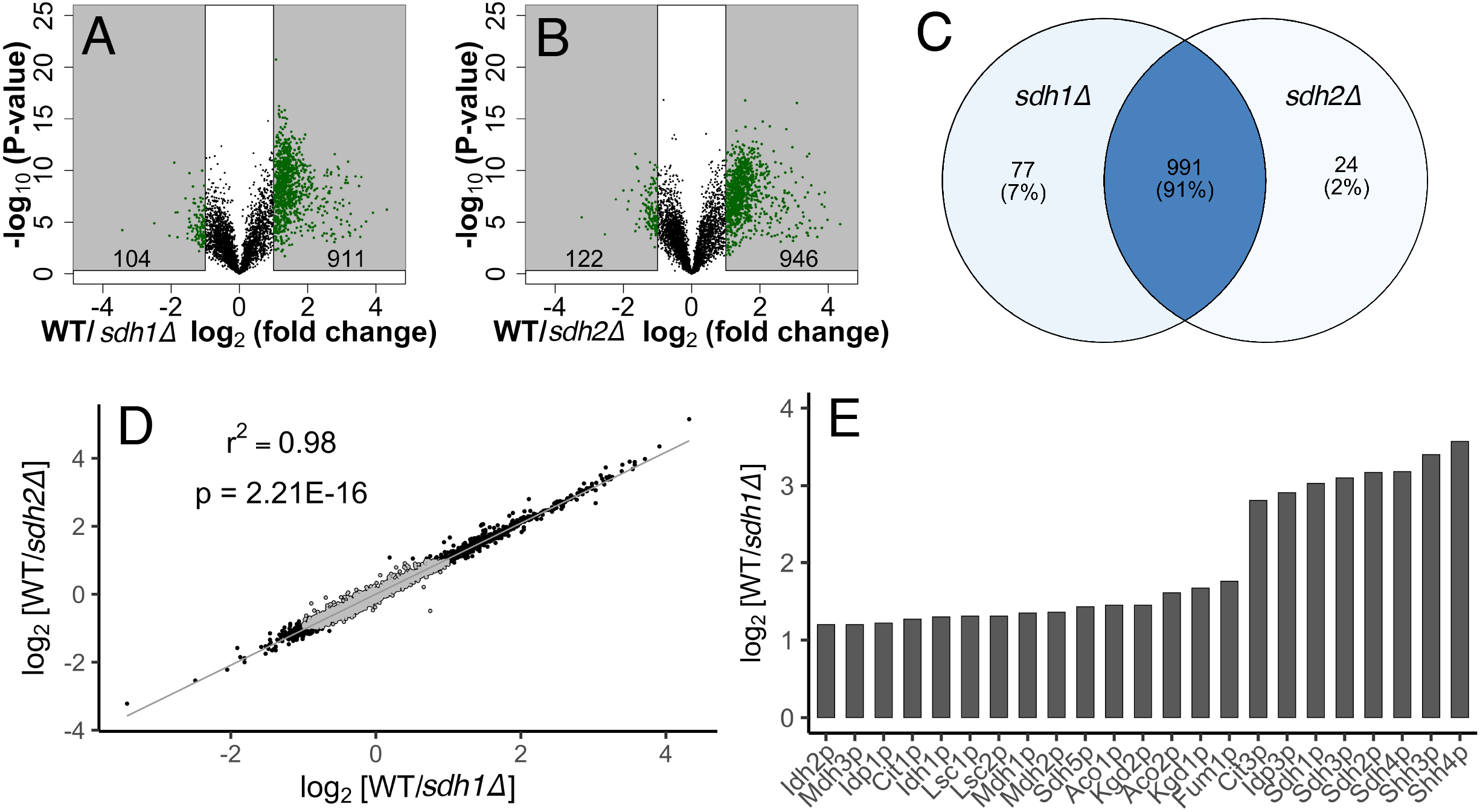
Proteomic comparison of *sdh1Δ* and *sdh2Δ* yeast strains. A, B. Volcano plots. Shading denotes significance (*p* < 0.05 and |log_2_ (fold change)| ≥ 1.5. Protein counts in each category are indicated. C. Venn diagram indicating relationship between proteins significantly different from WT for *sdh1*Δ and *sdh2*Δ *cells*. D. Correlation between *sdh1*Δ and *sdh2*Δ proteome alterations. Gray points indicate genes with non-significant fold changes. E. Results of DAVID functional analysis indicating loss of TCA cycle enzyme polypeptides, especially SDH subunits, in *sdh1*Δ yeast.

Applying pathway analysis using the DAVID Functional Annotation Tool (DAVID Bioinformatics Resources 6.8, NIAID/NIH), we found, as expected, that major proteomic changes in SDH-loss cells focused on enzymes of the TCA cycle. Quantitation of these enzyme changes is shown in (Fig. 3E). In this case, ratios are rough estimates due to peptide counting methods, such that complete absence of Sdh1p in *sdh1*Δ compared to WT yields a nominal ~10-fold change. Interestingly, loss of Sdh1p reduced most TCA cycle enzymes by 2-3-fold, but reduced SDH subunits (including Shh3p and Shh4p) to a greater extent.

The current proteomic data also allowed verification of the prior finding that both Sdh1p and Sdh2p are lost in *sdh1*Δ yeast, but a fraction of Sdh1p remains detectable in *sdh2*Δ yeast (Smith et al., 2007). Such evidence is shown in Fig. S4. This result is reminiscent of reports from mammalian cells where a complex of the SDHA catalytic subunit and chaperones appears to persist in the absence of SDHB (Bezawork-Geleta et al., 2018). Though there is no evidence of enzyme activity for the residual catalytic subunit, it might contribute in subtle ways to the different phenotypes of tumors driven by loss of different SDH subunits (Guzy et al., 2008). Thus, the small proteomic differences between *sdh1*Δ and *sdh2*Δ yeast may serve as a paradigm for small but important differences in comparable mammalian mutants affecting different SDH subunits. For example, the yeast Sdh3p subunit also functions as a component of the TIM22 mitochondrial translocase system, and yeast also expresses Shh3p and Shh4p paralogs of Sdh3p and Sdh4p, but with unknown functions (Gebert et al., 2011). The overall strong phenotypic and proteomic similarity of *sdh1*Δ and *sdh2*Δ strains led us to select the *sdh1*Δ strain as representative and appropriate for chemical suppression screening.

### LOPAC suppression screen

High-throughput screening (HTS) is an essential approach in drug discovery research (Macarron et al., 2011). We previously conducted a lethality screen of more than 200,000 compounds seeking agents selectively toxic to SDH-loss yeast cells as models of SDH-loss human familial PGL (Bancos et al., 2013). The current suppression screen focuses on the concept that the fundamental pathologies of SDH-loss cells are driven by the accumulation of succinate as an oncometabolite inhibiting 2-ketoglutarate-dependent dioxygenases (Cervera et al., 2009; Koivunen et al., 2007; Letouzé et al., 2013; Xiao et al., 2012). We hypothesized that ameliorating succinate toxicity may normalize cell function. Beyond succinate intoxication, it has been proposed that ROS also accumulate in SDH-loss cells, potentially compromising dioxygenase function by oxidation of the ferrous ion critical to dioxygenase function (Liu et al., 2020). The present suppression screen therefore was developed to identify compounds that selectively rescue growth of *sdh1*Δ yeast by restoring Jlp1p function using growth in ISE as the selection.

The LOPAC_1280_ library (Sigma-Aldrich) was screened at 10 μM in sixteen 96-well plates with 80 compounds per plate, allowing for growth controls (WT and *sdh1*Δ strains in AS and ISE media). Each experimental well was seeded with *sdh1*Δ yeast in ISE media and OD_600_ readings were taken at 7 timepoints over a 24-h period and a score modeled on the conventional Z statistic was calculated for each treatment and time (materials and methods). Compounds characterized by Z > 0 for at least 6 of 7 timepoints were classified as hits (30 of 1280 compounds; Fig. 4A).

**Fig. 4.**
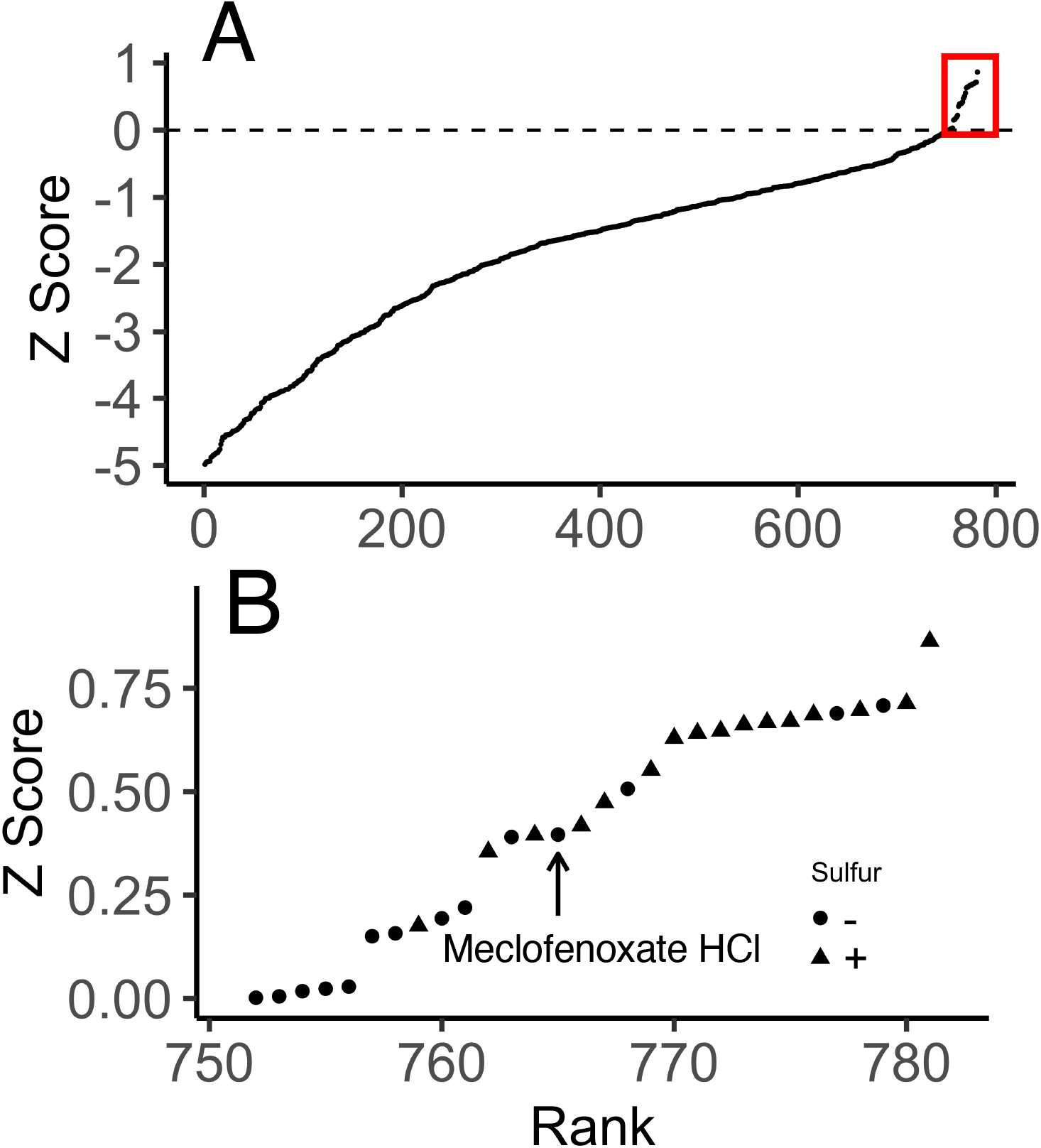
Results of LOPAC chemical suppression screen. A. Effects of LOPAC compounds on *sdh1*Δ yeast growth in ISE medium, ranked for compounds with Z Scores ≥ −5. Red box at upper right indicates compounds with Z ≥ 0. B. Detail of LOPAC compounds inducing Z Scores > 0, indicating whether the compound formulation itself did (+) or did not (−) contain bioavailable sulfate. Arrow indicates meclofenoxate HCl (Z = 0.40).

### Hit validation

Because the suppression screen was based on sulfur scavenging, drugs formulated with sulfate or related compounds (16/30 hits) were either excluded or more rigorously tested (Fig. 4B). Four of the initial 30 hits formed colored solutions that interfered with OD_600_ readings so were excluded. The seven compounds with highest z-scores and no confounding concerns (phenanthroline, phentolamine, reserpine, protoporphyrin IX, minocycline, pyrrolidinedithiocarbamate, and meclofenoxate) were repurchased and subjected to further validation (Supplemental Fig. S7). Validation assays were performed in 96-well plates grown without shaking. Oxygenation was judged adequate by the observation that the oxygen-dependent Jlp1p doxygenase in WT cells allowed strong growth in minimal ISE galactose media. Validation screening compared concentration-dependent effects of test compounds on the growth of WT, *sdh1*Δ, *sdh2*Δ and *jlp1*Δ strains on both ISE and AS media, seeking compounds that selectively improved growth only of *sdh1*Δ and *sdh2*Δ strains and only on ISE media.

This procedure led to the identification of meclofenoxate HCl (*The Science of Anti-aging Medicine*, 2003) as the most robust and reproducible hit (Fig. 5A). Meclofenoxate is an ester reported to rapidly hydrolyze in water to its substituents, dimethylaminoethanol (DMAE) and 4-chlorophenoxyacetic acid [Supplemental Fig. S2; (Yoshioka et al., 1987)]. Testing of these substituents showed DMAE to be the active agent (Fig. 5B, Supplemental Fig. S8). Meclofenoxate HCl and DMAE (25 μM) were then validated in a full comparative growth assay (Fig. 6), demonstrating their ability to selectively suppress the growth defect of SDH-loss yeast on minimal galactose medium with ISE as sulfur source. Selective partial suppression of the growth defect of SDH-loss yeast in ISE medium, without growth stimulation of WT or *jlp1*Δ strains in ISE media, or any of the strains in AS media, suggests that meclofenoxate HCl and DMAE act by improving Jlp1p catalysis of ISE conversion to sulfite. We sought evidence for potential mechanisms of this effect.

**Fig. 5.**
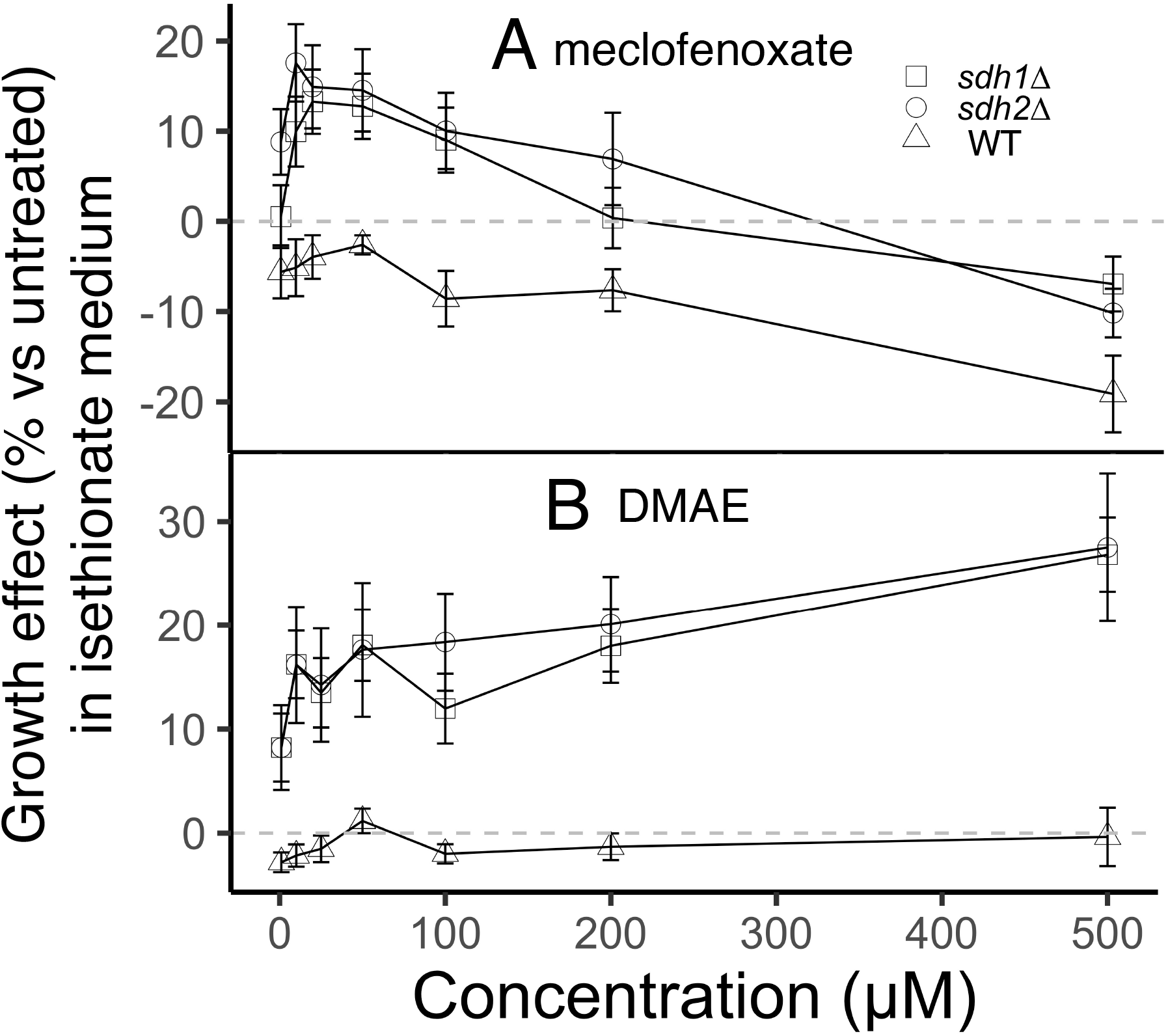
Dose-response data (percent change relative to untreated) for the indicated yeast strains in ISE medium and the indicated concentrations of (A) meclofenoxate and (B) DMAE, both dissolved in water. Error bars indicate standard deviation reflecting 4 technical replicates propagated to % growth effect vs. untreated.

**Fig. 6.**
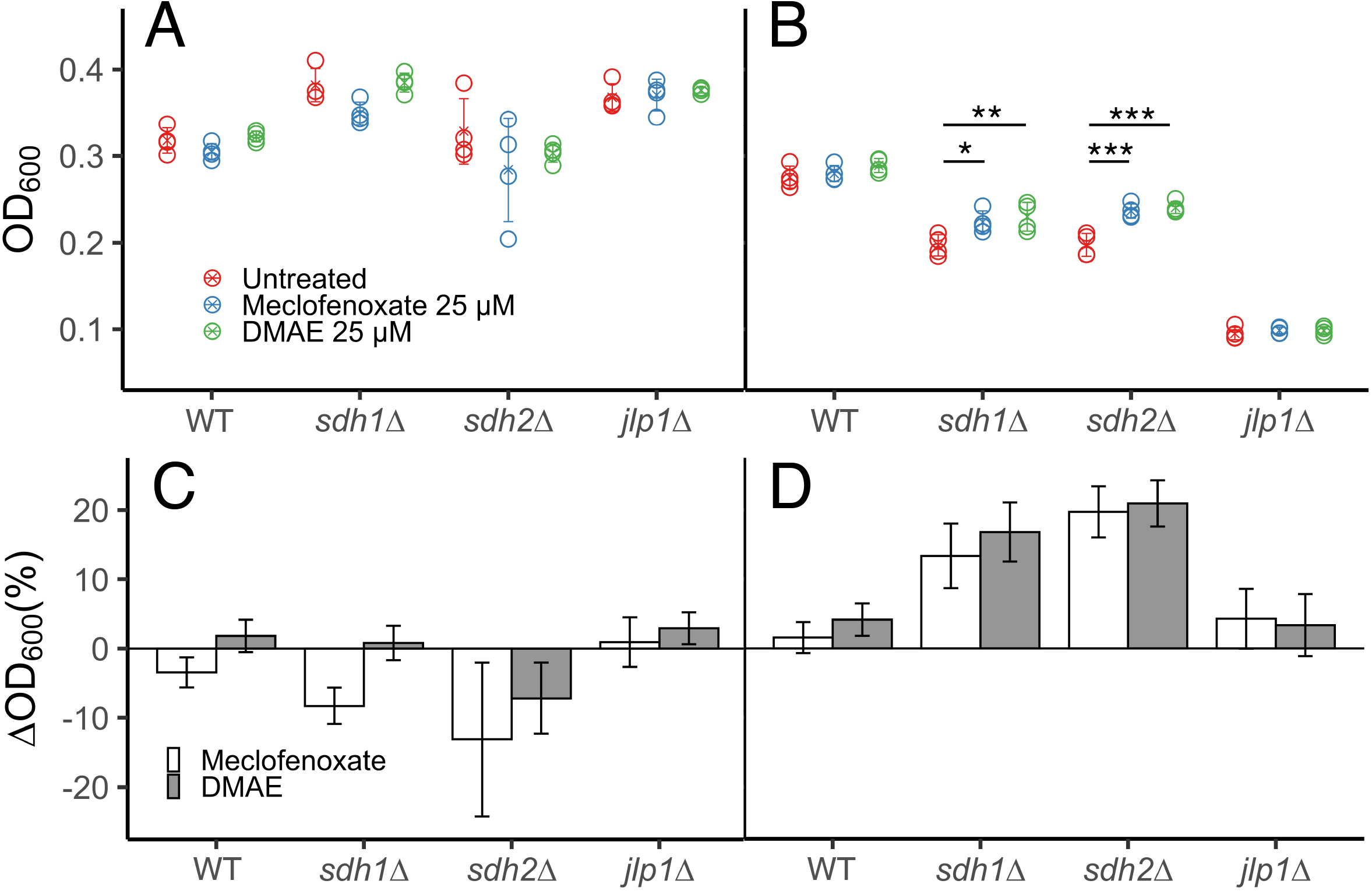
Effect of 25 μM treatment of the indicated drugs on yeast growth at 24 h in (A) AS and (B) ISE media. Drug effects are shown as % growth change in (C) AS and (D) ISE media for the indicated drug treatments. Statistical significance is reported using two-way ANOVA with a post-hoc Tukey HSD test for significance. *p*-values: * ≤ 0.05, ** ≤ 0.01, *** ≤ 0.001. In panel B Error bars indicate standard deviation of 4 technical replicates propagated to % growth effect vs. untreated.

### Metabolomics

Meclofenoxate HCl and DMAE have been studied as anti-aging treatments in animals and in assays of improved brain function (Dowson, 1985; Goldenberg, 1969; Kovalev et al., 2008; Marcer and Hopkins, 1977; Petkov et al., 1990; Zuckerman and Barrett, 1978; Miyazaki et al., 1976), but with little mechanistic detail. There is early published evidence that DMAE activates certain enzymes, including glucose-6-phosphate dehydrogenase (Bielenberg et al., 1986). We used NMR to monitor intracellular levels of succinate and 2KG, the metabolites whose concentration ratio is thought to determine dioxygenase inhibition (Fig. 7 AB). Remarkably, DMAE treatment significantly reduced intracellular succinate concentrations in both *sdh1*Δ and *sdh2*Δ yeast strains, but not in WT yeast. DMAE treatment also significantly increased 2KG levels in *sdh1*Δ yeast, but not in WT yeast. The resulting reduced succinate:2KG ratios are shown in Fig. 7C, providing evidence that DMAE treatment improves Jlp1p dioxygenase function by partially normalizing the intracellular succinate:2KG ratio in SDH-loss yeast cells.

**Fig 7.**
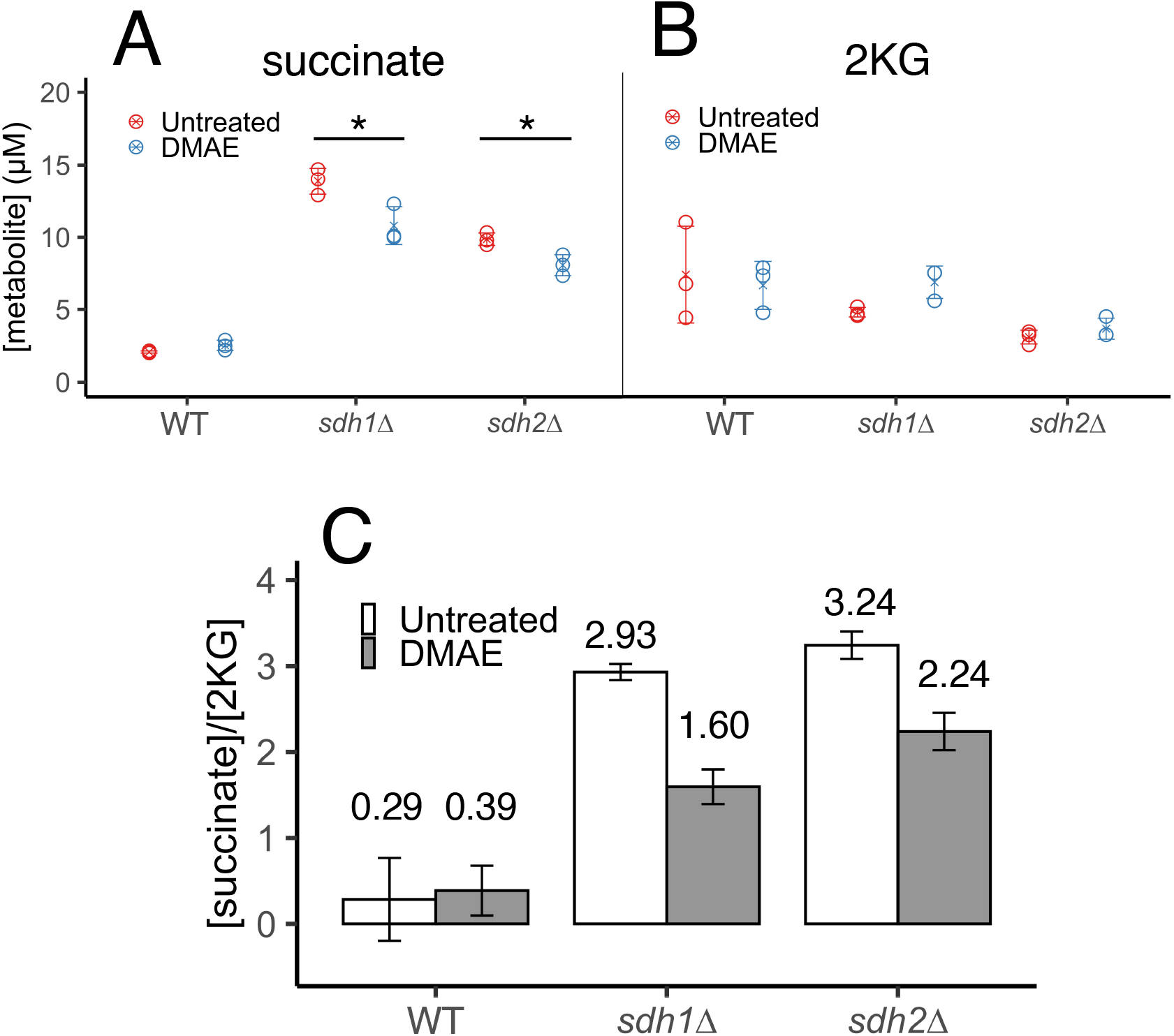
Effect of 100 μM DMAE on (A) intracellular succinate, and (B) intracellular 2KG concentrations in ISE medium after 24 h. Indicated level of statistical significance from a two-way ANOVA with a post-hoc Tukey HSD test for significance. * indicates P ≤ 0.05 based on 3 replicates. C. Ratio of succinate concentration to 2KG concentration in samples. Error bars represent standard deviation for 3 replicates.

### ROS assays

Though succinate toxicity is thought to be the primary oncometabolite in SDH-loss PGL, evidence shows that reactive oxygen species (ROS) may also accumulate and contribute to development of a cancer phenotype (Ishii et al., 2005; Smith et al., 2007). An altered intracellular redox state has the potential to shift the Fe^2+^: Fe^3+^ balance required for cycles of dioxygenase activity (Guzy et al., 2008; Liu et al., 2020). SDH-loss yeast have been shown to suffer increased ROS production with some increase in formation of protein carbonyl damage (Weber et al., 2015) but no acute evidence of toxic DNA damage (Smith et al., 2007). Assays monitoring the effects of DMAE treatment on protein oxidative damage yielded mixed results (Fig. 8A). DMAE treatment significantly reduced protein carbonyl levels in *sdh1*Δ cells but not in *sdh2*Δ or WT cells.

**Fig. 8.**
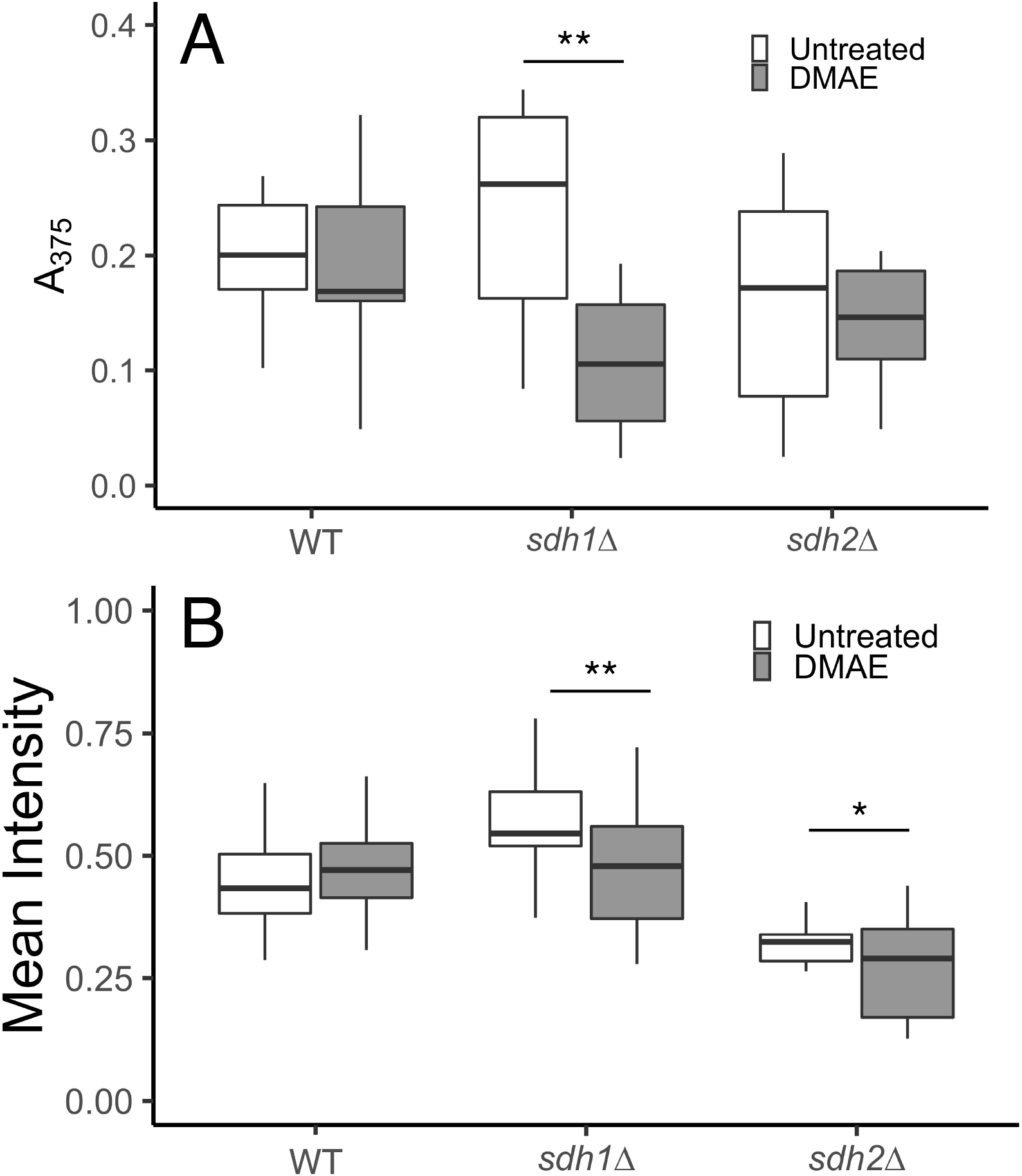
Effect of 100 μM DMAE treatment on measures of oxidative stress in ISE medium after 24 h. A. Assay of protein carbonyl products. B. Assay of ROS in live cells. Indicated level of statistical significance from a two-way ANOVA with a post-hoc Tukey HSD test for significance. * indicates P ≤ 0.05 and ** indicates P ≤ 0.01 based on 12 replicates (A) and 30-60 cells (B).

To more directly monitor ROS production, live cells were imaged via fluorescence microscopy after treatment with a fluorescein derivative activated by ROS. Results are shown in Fig. 8B and supplementary Fig. S5. Both *sdh1*Δ and *sdh2*Δ show significant decreases in fluorescence after treatment with DMAE while WT showed no effect. Interestingly, untreated *sdh2*Δ cells demonstrated a lower mean fluorescence intensity than untreated WT cells, different from earlier findings that ROS production was similarly increased in both *sdh1*Δ and *sdh2*Δ yeast (Smith et al., 2007). Replicative experiments were performed with dihydroethidine (DHE) staining. DMAE exhibited similarly subtle anti-oxidant activity (supplementary Fig. S6). These results suggest that anti-oxidant effects could also play a role in DMAE suppression of Jlp1p inhibition. Previous studies of such effects for DMAE have been limited to much higher concentrations (Gabriela Malanga et al., 2012).

### Preliminary DMAE testing in mammalian cells

The striking ability of meclofenoxate and DMAE to rescue a dioxygenase enzyme in SDH-loss yeast raises the obvious question of application in mammalian cells. Preliminary studies of up to millimolar concentrations of DMAE in WT and SDH-loss mammalian cells did not show normalizing effects on the intracellular succinate:2KG ratio (data not shown). Thus, while DMAE clearly supports the novel concept of suppressing succinate intoxication by altering metabolism in yeast, different compounds will be required for this purpose in mammalian cells.

### Implications

The disease consequences of metabolic perturbation are of central importance in medicine. Metabolic changes in cancer have been detected and discussed since the discovery of the Warburg Effect (de Alteriis et al., 2018; Kozal et al., 2021; Pavlova and Thompson, 2016; Warburg, 1956). SDH-loss familial PGL presents a unique metabolic state with its fragmented TCA cycle. The highly conserved nature of SDH across evolution makes our detailed proteomic comparison of *sdh1*Δ and *sdh2*Δ strains particularly interesting and relevant for understanding the long-standing paradox that, contrary to expectation for a multi-subunit enzyme, SDH-loss tumors show phenotypes dependent on the affected SDH subunit (Andrews et al., 2018; Guzy et al., 2008; Neumann et al., 2004; Rijken et al., 2019). While loss of any SDH subunit might be predicted to be equally disturbing to cell metabolism, there is evidence from human familial PGL that this is not true. By analogy with the mammalian case where residual SDH complexes may differ upon loss of certain subunits (Bezawork-Geleta et al., 2018), it is intriguing that SDH-loss yeast lacking Sdh1p show subtle proteomic differences from strains lacking Sdh2p. Rather than stressing the similarities between *sdh1*Δ and *sdh2*Δ for the purposes of our yeast screen, the subtle differences between the two strains may illuminate the differential penetrance of PGL caused by SDHA loss vs. SDHB loss in humans. There are ~20 differentially-expressed proteins between *sdh1*Δ and *sdh2*Δ yeast strains, and two interesting cases are COX19p and COX16p, both assembly factors for cytochrome c oxidase. These proteins are less abundant in the *sdh2*Δ strain than *sdh1*Δ strain, giving us a clue as to subtle proteomic differences that might have larger differences on disease progression among different human SDH-loss PGLs.

Research in familial PGL has focused primarily on three potential oncogenic mechanisms upon SDH loss: succinate accumulation driving dioxygenase inhibition (Koivunen et al., 2007; Selak et al., 2005; Smith et al., 2007; Xiao et al., 2012), ROS overproduction with corresponding damage and redox imbalance (Her and Maher, 2015; Ishii et al., 2005; Kregiel, 2012; Liu et al., 2020; Saffi et al., 2006), and succinylation of proteins leading to dysfunction (Smestad et al., 2018, 2017). Using the linkage between SDH loss and dioxygenase inhibition as a paradigm, the present chemical suppression screen exploited our ability to connect the inhibition of sulfur scavenging dioxygenase Jlp1p to a nutritional growth assay in a sulfur source requiring Jlp1p function in *S. cerevisiae* (Hogan et al., 1999; Smith et al., 2007). The leading hit from this screen, meclofenoxate HCl, and its active derivative DMAE, selectively increase growth of SDH-loss yeast strains by 10-20% in ISE media.

Though studied superficially over many years, potential mechanisms of meclofenoxate and DMAE effects on metabolism, physiology and lifespan remain poorly understood. Initial studies in the 1970s suggested that these drugs substantially enhance the lifespan of mice (Hochschild, 1973; Miyazaki et al., 1976). DMAE was found to affect brain tissue, reducing levels of lipofuscin, an oxidative plaque associated with aging. Lohr and Acara noted that DMAE is similar in structure to choline and could potentially serve as a precursor, and that it could also inhibit choline oxidase (Lohr and Acara, 1990), reducing levels of betaine. Anti-aging effects of DMAE have been studied with an eye to a potential free-radical scavenging mechanism (Gabriela Malanga et al., 2012).

Perhaps the most tantalizing published mechanistic observation for DMAE is its reported ability to increase the activity of glucose-6-phosphate dehydrogenase, the rate-limiting initial enzyme of the Pentose Phosphate Pathway (PPP) (Roy and Singh, 1983). The PPP runs parallel to glycolysis, but generates reducing equivalents and key carbon skeletons for nucleotide biosynthesis without generating ATP. We hypothesize that agents such as DMAE may enhance dioxygenase function in SDH-loss cells by shunting carbon flux away from glycolysis and the TCA cycle, reducing the production of succinate at the SDH blockade. We provide further evidence that DMAE relieves oxidative stress by reducing ROS, perhaps normalizing the obligatory Fe^2+^ - Fe^3+^ equilibrium essential to the dioxygenase catalytic cycle.

Thus, this work illustrates a new paradigm for rewiring metabolism through a small molecule cue that reduces oncometabolite accumulation. Searching for effective small molecules of this type that might affect SDH-loss PGL tumor cells offers a potential route to normalizing inhibited dioxygenase function in such cells. Because dioxygenase inhibition by succinate causes both pseudohypoxia and histone and DNA hypermethylation, such metabolic rewiring could have broad therapeutic effects in familial PGL.

## Funding

This work was supported by the Mayo Foundation, the Mayo Clinic Graduate School of Biomedical Sciences, and the Paradifference Foundation.

## Author contributions

Project conception and manuscript preparation: WB, MB, LJM

Experiments: WB, MB, KS, KP, BW

Data analysis: WB, MB, KS, KP, BW

## Acknowledgments

We thank members of the Maher laboratory for assistance and Marina Ramirez-Alvarado for sharing instrumentation. We acknowledge the excellent assistance of members of the Katzmann laboratory at Mayo Clinic, especially Chun Che Tseng. The excellent support of Ivan Vuckovic and Song Zhang in the Mayo Clinic NMR core facility, Akhilesh Pandey, Zachary Ryan, and Benjamin Madden in the Mayo Clinic Proteomics core facility, and Mai Petterson and Ian Lanza in the Mayo Clinic Metabolomics core facility are acknowledged.

## Notes

### Competing Interest Statement

The authors have declared no competing interest.

