## Supplemental Materials for "A chemical screen for suppressors of dioxygenase inhibition in a yeast model of familial paraganglioma"

and L. James Maher, III*

### Supplemental methods

#### Yeast genotyping

Yeast cells were subjected to centrifugation at 2500 × *g* and the pellet was washed in DTT buffer (100 mM Tris pH 9.4, 10 mM DTT), then resuspended in 100 µL zymolyase buffer [1 M sorbitol, 20 mM Tris pH 7.5, 50 mM EDTA, 1% β-mercaptoethanol, 1-2 mg/mL Zymolyase (AMSBIO #120493-1)] for cell wall lysis at 30˚C for 15 min. Quenching buffer (400 µL: 0.5% SDS, 100 mM Tris pH 8.0, 50 mM EDTA) was added and incubated at 70˚ C for 15 min. Potassium acetate solution (5 M, 100 µL) was added and incubated on ice for 15 min. The resulting mixture was subjected to centrifugation at 14,000 × *g* and the supernatant decanted into a fresh tube. Ethanol (900 µL) was added with incubation for 5 min at room temp. Cells were again pelleted, washed once with 70% ethanol, resuspended in 1 mL water, and DNA quantified using a Nanodrop spectrophotometer. Genomic DNA was amplified using PCR. Conditions were 5 mM $MgCl_2$, 1× PCR Buffer (ThermoFisher), 0.1 mg/mL BSA, 0.2 mM dNTPs, 0.625 U/25 µL Taq polymerase, 1 µM each primer, with 50 ng template). the PCR cycle was: 1× 94 ˚C for 3 min, 35× [94˚C for 15 sec; 57 ˚C for 15 sec; 72 ˚C for 1 min], 1× 72 ˚C for 3 min, with subsequent cooling. Primers were obtained from IDT (Coralville, IA). PCR products were subjected to electrophoresis through a 1.5% agarose gel in TAE buffer for 60 min and imaged with an Amersham Typhoon Instrument. Resulting gel images were analyzed using FiJi.

**Metabolite analysis**

500 µL of each sample (original growth media and cell lysate) were transferred to NMR tubes and mixed with 150 µL NMR buffer (100 µL phosphate buffer and 50 µL TSP/$D_2O$). TSP/$D_2O$ is 99.9% $D_2O$ and 0.05% (w/v) TSP as an NMR indicator. Intracellular and extracellular metabolite concentrations were determined using quantitative NMR spectroscopy. The $^1$H NMR spectra were acquired using a 600 MHz spectrometer (Bruker BioSpin, Billerica MA) equipped with a 5 mm BBI room temperature probe and an automatic refrigerated sample changer (SampleJet). The sample temperature in the magnet was regulated to 298.2 ± 0.1 K with a BTO 2000 variable temperature unit. The spectra were recorded with water peak suppression using the standard 1D NOESY pulse sequence (noesygppr1d; Bruker BioSpin), acquiring 128 scans, with 65,536 data points, a spectral width of 12,019 Hz, a mixing time of 10 ms and a relaxation delay of 4 s. The phase and baseline corrections were performed using TopSpin v3.6 (Bruker Biospin). Metabolites were identified and quantified using the software program Chenomx NMR Suite v8.5 (Chenomx, Edmonton CA), by fitting the spectral lines of library compounds into the recorded NMR spectrum of the cell extracts. The quantification was based on peak area of TSP-$d_4$ signal.

**Proteomic analysis**

Frozen mitochondria were thawed and lysed with modified RIPAA buffer (5% SDS, 1% NP-40, 0.5% sodium deoxycholate, 1 mM EDTA, 50 mM Tris pH 8.0, protease inhibitor cocktail [Roche 11836153001], phosphatase inhibitor cocktail [ThermoFisher 78442]). Proteins were clarified by centrifugation and protein concentrations were determined using a BCA assay kit according to manufacturer's instructions (ThermoFisher 23227). Proteins were then digested to peptides using an S-Trap mini according to manufacturers instructions (Protifi). S-trap eluted peptide digests were quantified in a peptide assay (ThermoFisher 23275). Peptides were then dried, reconstituted in 100 mM triethylammonium bicarbonate buffer (pH 8.5) and labeled with 500 mg of the TMT 10-plex isobaric reagents (Thermo Fisher Scientific, Waltham, MA, LOT# UK291565) following the manufacturer's instructions with 1 h reaction at room temperature and quenching with 5% hydroxylamine. Excess TMT reagents were removed from the pooled mixture using $C_{18}$ Sep Pak Plus cartridges (Waters, Milford MA), lyophilized and fractionated using basic pH reverse phase HPLC on a Thermo Ultimate 3000 RSLC HPLC system with a Waters XBridge peptide BEH C18, 3.5 µm, 4.6 mm × 250 mm column, running a 5% B to 60% B gradient over 60 min with 5 mM ammonium formate as the A solvent and 5 mM ammonium formate/90% acetonitrile as the B solvent. 96 fractions were collected over the 80-min run and were concatenated to 24 fractions for nanoLC-tandem orbitrap mass spectrometry analysis on an Orbitrap Exploris 480 mass spectrometer (Thermo Fisher Scientific, Waltham, MA) coupled to an Ultimate 3000 RSLCnano system (Thermo Fisher Scientific, Waltham, MA). Peptides were loaded onto a trap column (EXP2 Stem Trap, 2.7 µm, Halo $C_{18}$, 0.18 × 13.5 mm; Optimize Technologies) and separated on an analytical column (EasySpray, $C_{18}$ 75 mm × 50 cm; Thermo Fisher Scientific). Mobile phase A and mobile phase B were composed of 0.2% formic acid in 98% water/2% acetonitrile and 0.2% formic acid in 80% acetonitrile/10% isopropanol/10% water, respectively. Peptides were separated over 90 min from 5 to 35% of mobile phase B at a flow rate of 300 nL/min. The mass spectrometer was operated in a data-dependent mode with a cycle time of 3 s. MS1 scans (350-1600 m/z) were acquired with an orbitrap resolution of 120,000, automatic gain control (AGC) of 100% and maximum injection time of 50 ms. Precursor ions were isolated in the quadrupole with an isolation width of 0.7 m/z and fragmented with high-energy collision induced dissociation (HCD) at a normalized collision energy of 39%. MS/MS scans were acquired with an orbitrap resolution of 60,000, AGC of 200% and a maximum injection time of 105 ms. Raw data were searched against the SwissProt yeast database (release 2021_01, 6637 entries) using Sequest HT in Proteome Discover (ver 2.5. Thermo Scientific) with the parameter settings at 10 ppm for the precursor mass tolerance, 0.6 Da for the fragment mass tolerance and full trypsin specificity. Carbamidomethylation of cysteines and TMT label (+229.163) of peptide N-termini and lysines were set as static modifications and oxidation of methionine and acetylation of protein N-termini were considered as dynamic modifications during the database search. Identification was based on a 1% false discovery rate threshold applied at peptide and protein levels. TMT reporter ion quantitation was based on PSMs with a co-isolation threshold of <50% with an average reporter signal-to-noise threshold of at least 10. The abundance values were normalized to the same total peptide amount per channel and scaled so that the average abundance per protein and peptide was 100. Data analysis was performed using Microsoft Excel and R.

### Supplemental Table S1. Yeast minimal galactose medium components

| Component | Final concentration |
|---|---|
| Ammonium chloride | 15 mM |
| Potassium phosphate monobasic | 6.6 mM |
| Potassium phosphate dibasic trihydrate | 0.5 mM |
| Sodium chloride | 1.7 mM |
| Calcium chloride | 0.7 mM |
| Magnesium chloride hexahydrate | 2 mM |
| Boric acid | 0.5 µg/mL |
| Copper chloride (2 H2O) | 0.0447 µg/mL |
| Potassium iodide | 0.1 µg/mL |
| Zinc chloride | 0.19 µg/mL |
| Ferric chloride | 0.03 µg/mL |
| Calcium pantothenate | 2 µg/mL |
| Thiamine HCl | 2 µg/mL |
| pyridoxine | 2 µg/mL |
| inositol | 20 µg/mL |
| biotin | 0.02 µg/mL |
| Galactose | 2% |
| Ammonium sulfate OR Isethionate | 20 µM |
| Leucine | 0.06 mg/mL |
| Lysine | 0.075 mg/mL |
| Histidine | 0.04 mg/mL |
| Tryptophan | 0.04 mg/mL |
| Adenine | 0.04 mg/mL |
| Uracil | 0.04 mg/mL |

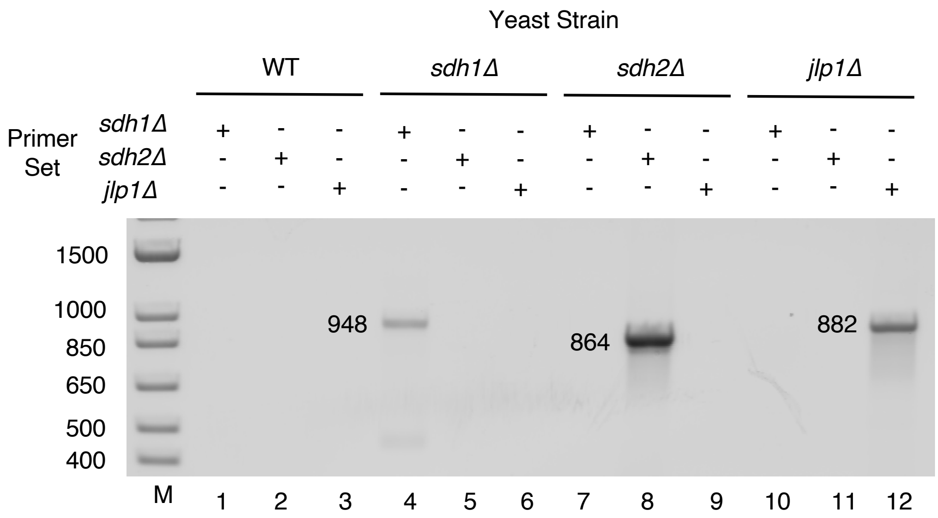


Fig. S1. Validation of yeast genotypes. PCR amplification of knockout genes of interest employed primers for each of the three knockout strains. Predicted PCR product lengths (bp) are indicated.

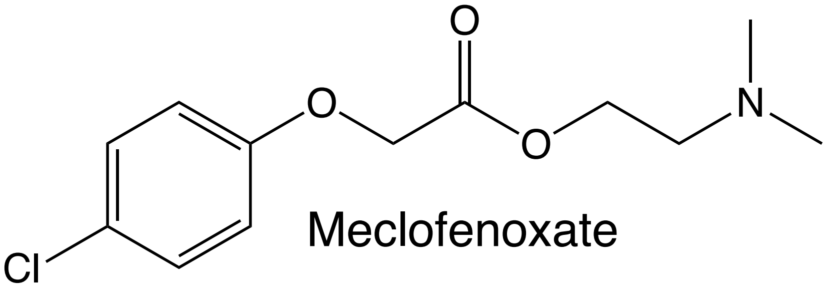

Meclofenoxate

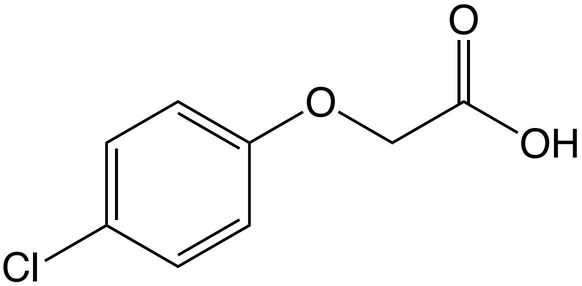

*p-chlorophenoxyacetic acid*
(4-CPA)

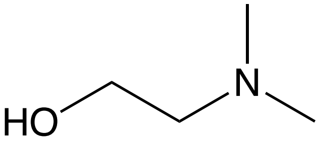

Dimethylaminoethanol
(DMAE)

Fig. S2. Structures of Meclofenoxate ester and its derivatives.

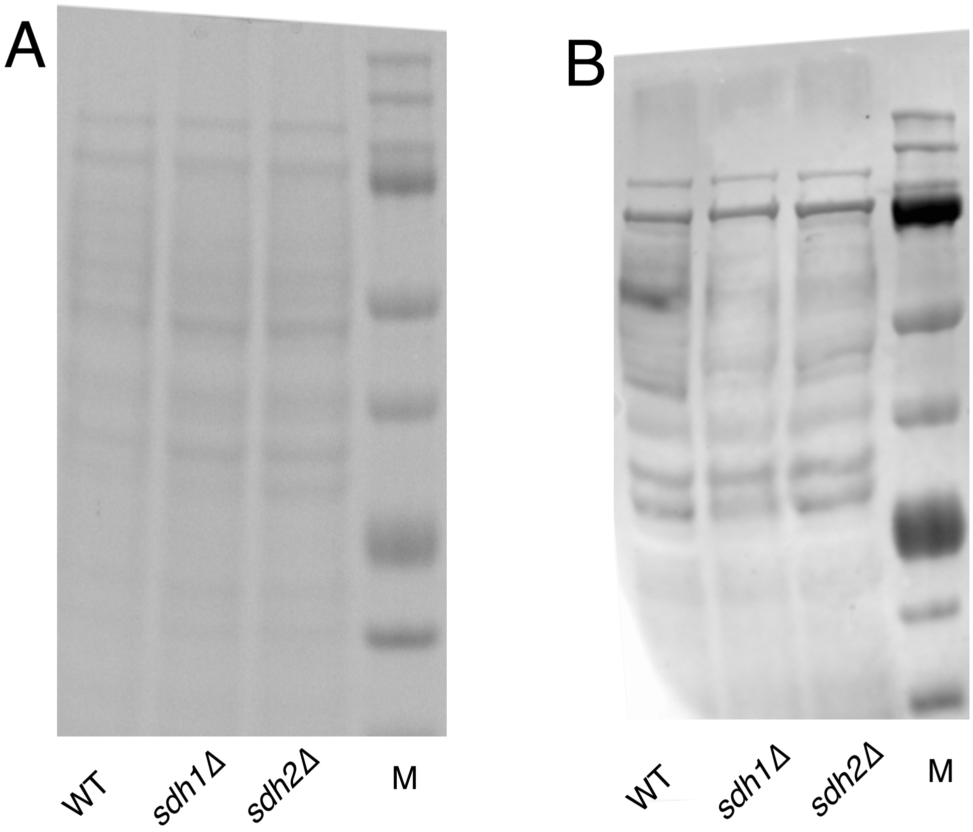


Fig. S3. Analysis of mitochondrial protein succinylation. A. Coomassie-stained gel showing equal protein loading. B. mitochondrial extracts from WT, *Sdh1*Δ, and *Sdh2*Δ yeast grown to saturation in YPGal. Western blot using anti-pan-succinyllysine antibody.

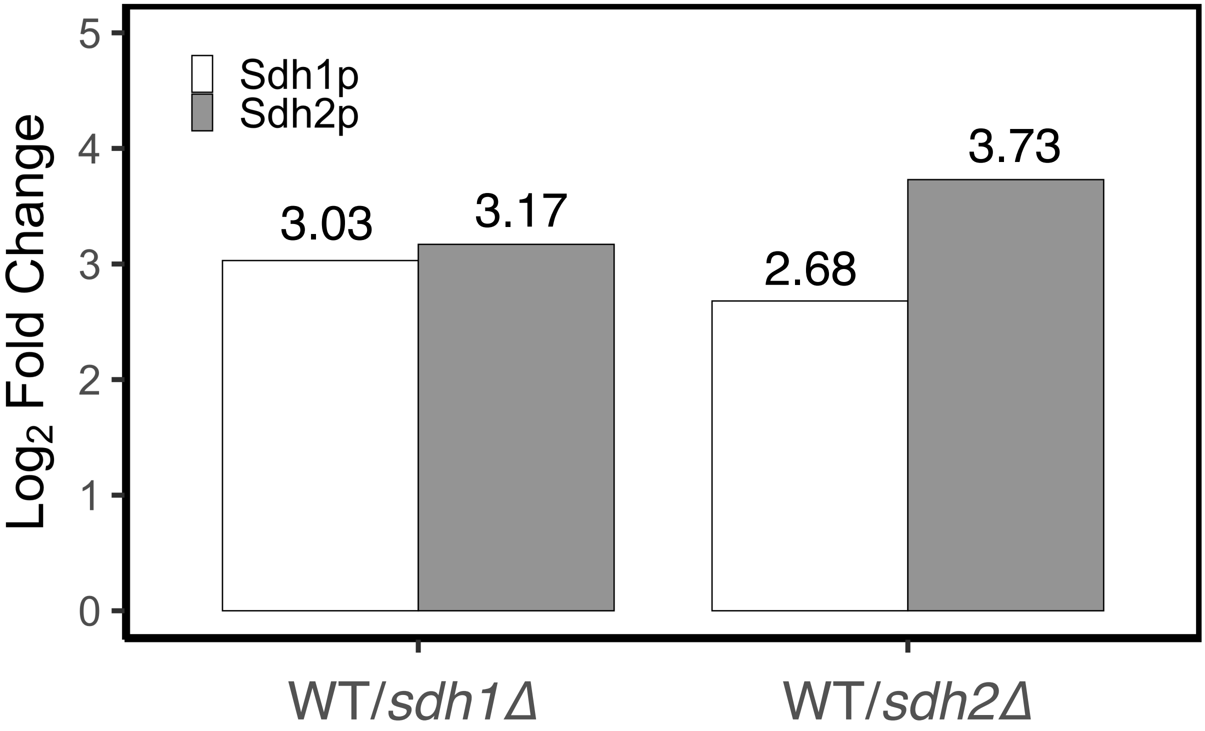


Fig. S4. Residual SDH subunit comparison.

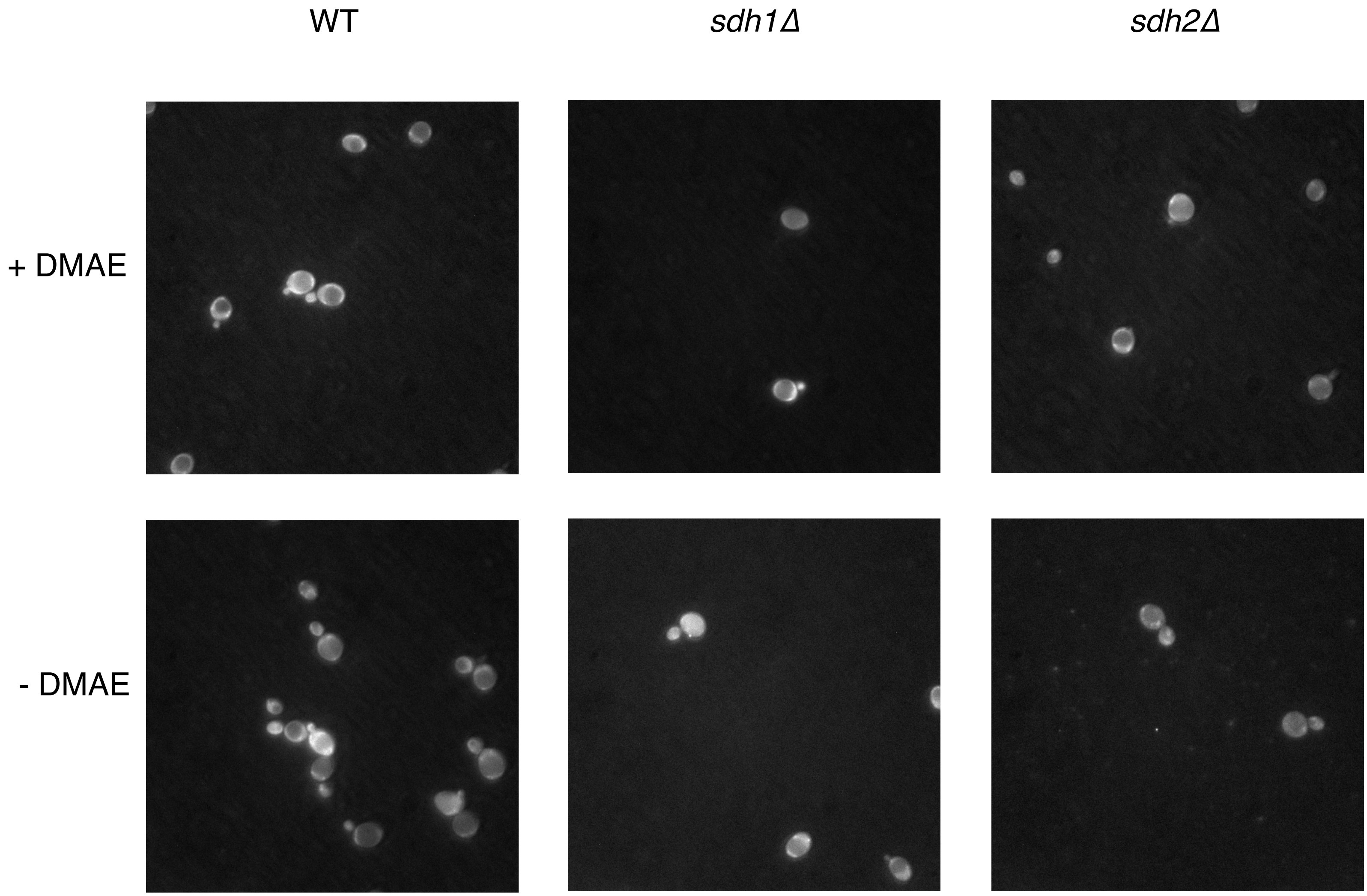


Fig. S5. Representative confocal microscopy images of yeast live cell ROS stained with $H_2$DCF-DA. Magnification 100 ×. Excitiation using FITC filter and emission using PE-TR filter.

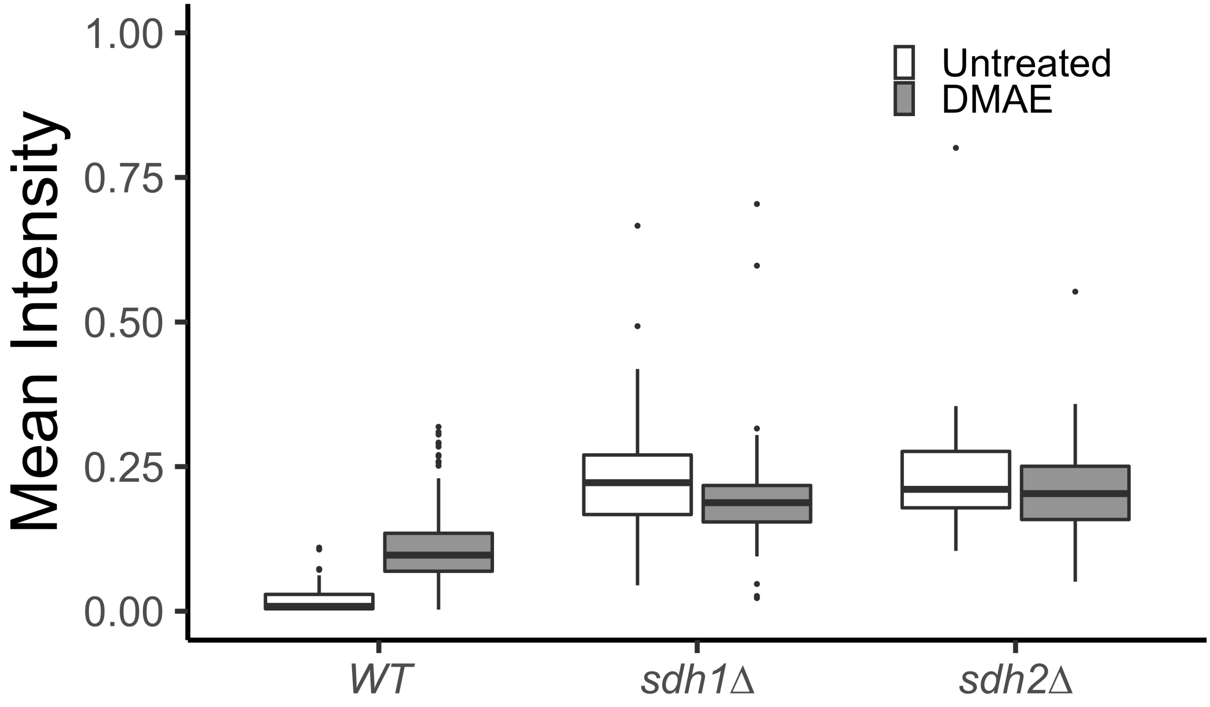


Fig. S6. Effect of 100 μM DMAE treatment on ROS in ISE medium after 24 h. Assay of ROS in live cells using Dihydroethidium (DHE). Indicated level of statistical significance from a two-way ANOVA with post-hoc Tukey HSD test for significance where * indicates $P \leq 0.05$ based on 30-60 cells.

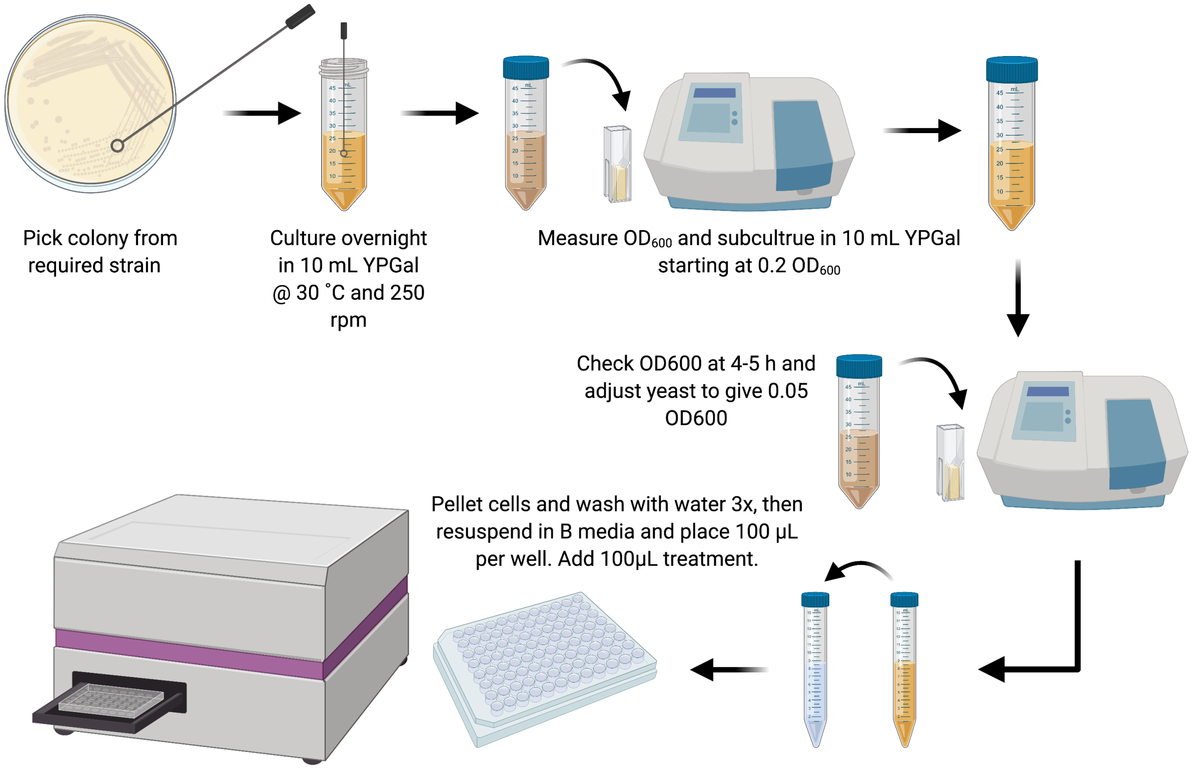


Fig. S7. Workflow of the yeast growth assay for testing drug candidates.

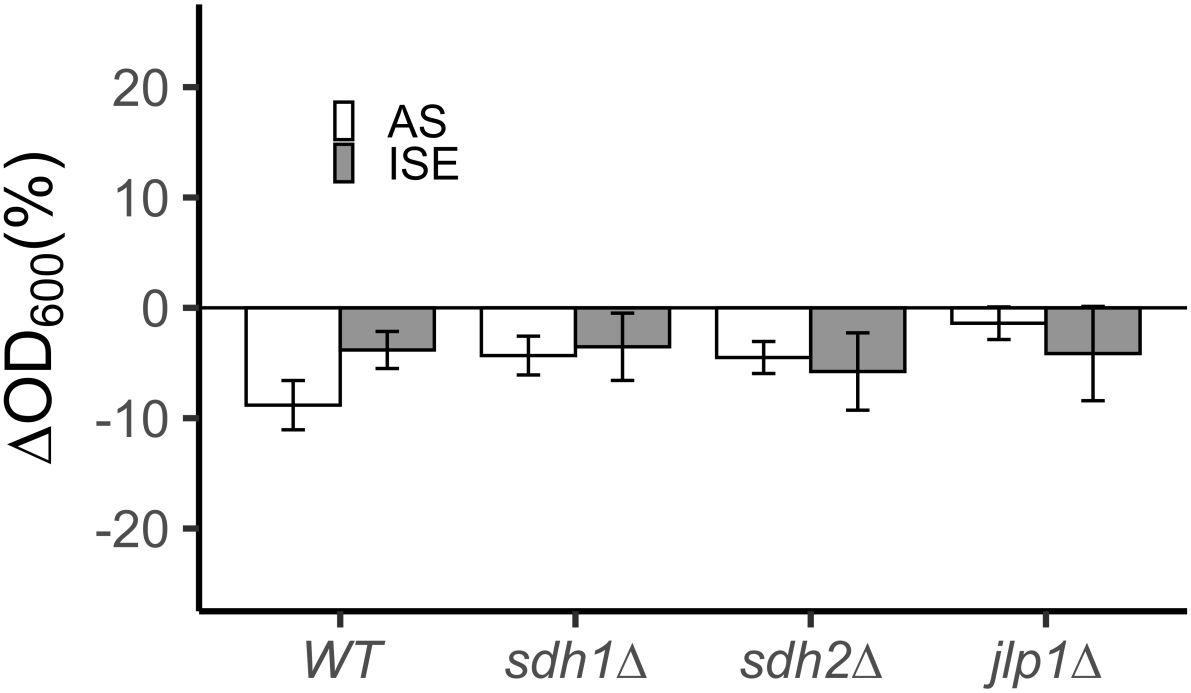


Fig. S8. Effect of 4-chlorophenoxyacetic acid (10 μM) on yeast growth at 24 h in AS and ISE media. Drug effects are shown as % growth change from untreated yeast in AS and ISE. Error bars indicate standard deviation of 4 technical replicates propagated to % growth effect vs. untreated.
